# Small stream restoration increases habitat and plant diversity across scales

**DOI:** 10.64898/2026.06.23.733988

**Authors:** Lena Lerbs, Alina Singer, Anna Dotzert, Lisa Grunwald, Jonas Lampe, Nina Farwig, Sascha Liepelt, Franziska Willems, Stefan Pinkert, Anna Bucharova

## Abstract

A majority of stream restoration efforts in central Europe focus on streams that are less than five meters wide. Restoration aims to increase structural complexity, thereby enhancing habitat heterogeneity, promoting biodiversity, and reestablishing aquatic-terrestrial linkages that can drive responses in adjacent terrestrial communities. However, the effects of small stream restoration on terrestrial biodiversity remain poorly understood because research focuses mainly on large rivers.

Here, we investigated the effects of restoration on the structural complexity of the stream channel as well as the terrestrial habitat and plant diversity on a local and landscape scale across 55 small streams in an agricultural landscape. We compared restored stream sections with non-restored ones that were similar to the conditions before restoration. We also assessed how restored sections changed over time since restoration.

Restored stream sections showed a higher stream structural complexity and habitat diversity, both of which are targets of active restoration measures. Restoration also increased riparian plant diversity, both directly and indirectly through structural complexity and habitat diversity. Although time since restoration did not influence structural complexity, it drove successional changes in plant communities that became increasingly associated with wetland habitat conditions.

Our results demonstrate that small stream restoration effectively increases floodplain habitat and plant diversity in agricultural landscapes, primarily by enhancing water availability in the floodplain. Restoration actions on small streams support biodiversity if they improve stream channel complexity, connect the stream with its floodplain, and create floodplain habitats.

## 1. Introduction

Riparian areas are biodiversity hotspots shaped by hydrological disturbance regimes, but humans have altered these ecosystems for centuries (Riis et al., 2020; Tockner & Stanford, 2002). Particularly, stream channelization and floodplain drainage for settlements, agriculture and forestry have profoundly altered floodplain dynamics (Naiman et al., 2013; Zerbe & Wiegleb, 2009). River modifications often caused the water table to drop by several meters, which disconnected the river from its floodplain and reduced both the frequency and duration of floods (Leuschner & Ellenberg, 2017). The absence of natural dynamics, combined with increased anthropogenic pressures from pollution and climate change, led to habitat degradation, homogenization and biodiversity loss at both local and landscape scales (Dudgeon et al., 2006). Degraded floodplains also provide reduced water retention capacity which contributes to catastrophic floods further downstream (Reid et al., 2019). An increasingly popular approach to counteract these negative effects is river and floodplain restoration.

Restoration projects on rivers and streams aim to improve water retention, river dynamics and biodiversity (Pilotto et al., 2019; Verdonschot & Verdonschot, 2023). Common restoration measures include hydro-morphological enhancements like re-meandering, raising the stream beds and adding boulders or large dead wood to improve the structural diversity of the stream (Kollmann et al., 2019; Morandi et al., 2017). These measures successfully alter hydrological conditions, create wetter habitats, improve flow dynamics and the number of microhabitats (Januschke et al., 2011; Modrak et al., 2017). While physical habitat conditions show consistent improvement, the response of aquatic taxa is weak to absent (Hering et al., 2015; Lorenz & Feld, 2013). Restoration of stream sections hardly improves the water quality that is needed for the recovery of aquatic biota, and unrestored sections hinder recolonization of the restored sections (Lepori et al., 2005; Muhar et al., 2016). This indicates that hydro-morphological restoration does not uniformly translate into recovery of aquatic communities.

However, river and stream restoration can have positive effects on adjacent terrestrial communities in the riparian zone because they are less constrained by water quality and dispersal limitations (Januschke et al., 2014; Wenskus et al., 2025). Additionally, restoration measures often include active establishment of diverse riparian vegetation (Fraaije et al., 2019; Morandi et al., 2017). Riparian vegetation provides important ecosystem functions such as habitats for other organisms, regulation of the microclimate and improving water quality through filtering pollutants and nutrients (Naiman & Décamps, 1997). As the riparian vegetation is tightly linked to the dynamic river system and is affected by occasional changes of the river bed and disturbance through floods (Baumane et al., 2024), it can serve as linkage from the stream to the terrestrial side of the stream. Recovery of riparian vegetation thus can be a direct consequence of improved physical conditions in restored streams and may indicate broader ecological improvements (Riis et al., 2020; Vannote et al., 1980).

The ecological effects of stream restoration develop over time as physical processes re-establish and reshape floodplain habitats. Frequent disturbances create a mosaic of vegetation types that is changing through time and is rarely in an equilibrium state (Ward et al., 1999). Floodplains are therefore characterized by high vegetational heterogeneity, and heterogeneous habitats typically host higher biodiversity following the habitat heterogeneity hypothesis (Stein et al., 2014; Tews et al., 2004). After successful restoration, riparian vegetation depends on the restored hydrodynamic processes (Hering et al., 2015; Modrak et al., 2017). However, the success is not immediate, because flood regimes need time to re-establish, and so does the associated formation of mosaics of floodplain habitats (Hasselquist et al., 2015; Lennox et al., 2011; Nilsson et al., 2015). The response of vegetation to river restoration thus changes through time.

Our understanding of restoration effects on riparian biodiversity comes primarily from restoration projects on large rivers. However, the majority of river restorations take place on streams that are less than five meters wide (Bernhardt et al., 2005; Morandi et al., 2017).

Findings from large rivers cannot be directly transferred to small streams, because small streams differ from larger rivers in their discharge, water chemistry and volume while they are particularly sensitive to pesticide and nutrient inputs from the surrounding landscape (Kristensen & Globevnik, 2014; Meyer et al., 2007). However, studies on the effect of small stream restoration on terrestrial vegetation are rare so far.

To address this gap, we investigated the effect of small stream restoration on the structural complexity of the stream as well as the habitat and plant diversity in the riparian area, particularly the presence of floodplain habitats and wetland plant species. We mapped habitat types and recorded plant species composition at 55 restored sections of small streams and compared them to paired unrestored controls. We hypothesize that (1) restoration has a positive effect on stream structural complexity, habitat richness, as well as on the occurrence of floodplain habitats. (2) Plant species richness (alpha diversity), and the dissimilarity of plant communities (beta diversity) will therefore increase on a local and landscape scale, while the moisture preference of plant species turns towards wetlands.

Consequently, restored and control sections host different plant communities. (3) The positive effects of restoration on plant diversity are mediated by increased structural complexity of the stream and habitat richness. (4) The positive effects of restoration on habitat and plant diversity increase with time since restoration.

## 2. Material and methods

### 2.1. Study design

We surveyed vegetation at 55 small low mountain range streams that were actively restored 2 to 25 years prior to the data collection. All streams belong to category III of the local river classification (Hessisches Wassergesetz, 2010), which is headwaters. They were in agricultural areas in central Germany, and shared similar climatic, landscape, and elevation contexts. The maximum distance between sites was 55 km (Fig. 1).

**Figure 1:**
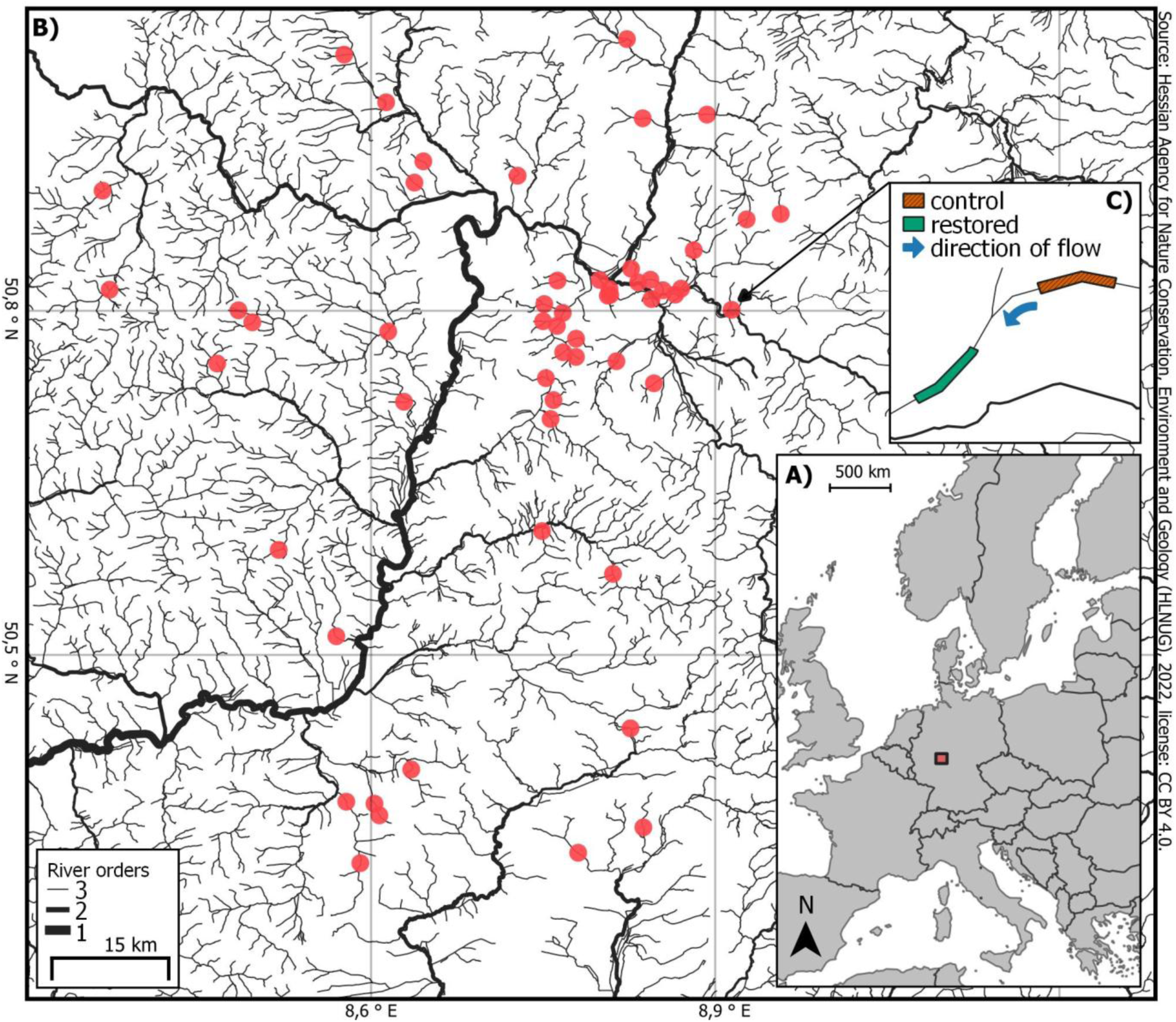
Study area. **A)** Location of the study area in central Europe. **B)** Overview of 55 study sites on small headwater streams. **C)** Exemplary placement of control and restored site along the stream direction.

We established a paired design to compare restored and non-restored, degraded (control) sections (Pickett, 1989). For each restored section, we determined project boundaries using available documentation and historic aerial photographs in Google Earth Pro (Google LLC., 2024). For each restored section, we selected a control section nearby that resembled the pre-restoration state of the restored section, based on pre-restoration aerial imagery. The control section matched its corresponding restored section by length, area and shape. In total, there are 110 sections, 55 restored and 55 corresponding non restored. The restored and its corresponding control section are further referred to as pair identity number.

Whenever possible, we placed the control sections upstream, to prevent a spill-over of restoration effects into the control section. When upstream placement was not possible, we placed the controls downstream (18%) or on a comparable stream nearby (24%). Restored and corresponding control sections were on average 0.4 km apart, with a maximum distance of 1.6 km. Selection and delineation of the sections was done in the geographic information system QGIS (QGIS Development Team, 2023).

### 2.2. Stream structural complexity

To evaluate the effect of restoration on stream morphology, and to differentiate between more and less intensive restoration measures, we assessed the stream structural complexity of both restored and control sections in 2025 for 53 out of 55 paired sections. To do this, we used the official sampling protocol of the German working group on water issues (LAWA, 1999; see summary of methodology in supplementary material). We assessed the stream structural complexity of 100 m reaches, with the number of reaches varying between 1 and 7 per section, depending on the length of the stream within the section. Along these reaches we then evaluated the complexity of the flow, the longitudinal and cross profile, the bed and bank structure, and calculated the stream’s structural complexity index (see modified calculation in supplementary material). We rescaled the original stream structural complexity index to range between 0 to 1 with higher values indicating higher stream structural complexity.

### 2.3. Habitat diversity

To investigate habitat diversity, we assessed the habitat richness and the proportion of key floodplain habitats in 53 paired sections. For this, we mapped and classified habitat types of the entire section based on phytosociological plant associations according to Schubert et al. (2010) in May and June 2025. Habitat types (hereinafter called “habitats”) represent ecologically coherent vegetational units defined by a characteristic species composition (see Table S4 for an overview of habitats). We calculated habitat richness of each section as the total number of distinct habitats present in each section. Of these habitats, we identified key floodplain habitats using the translation table provided by Januschke et al. (2024, see Table S4) and calculated the proportion of area they cover per section.

### 2.4. Plant diversity

We recorded plant species and their cover in 450 plots of 4 m × 4 m size in all 55 paired sections. The number of plots per section was proportional to the section area, specifically, it increased with square root of the section area and ranged between two and nine per section. Before fieldwork, we used aerial imagery to place the plots within the sections to cover all visible habitats. As paired restored and control sections had equal section areas, the number of plots was identical within the pair.

We surveyed vegetation on 198 plots during May and June 2023 and on 252 plots in May and June 2024, with the restored and respective control section being surveyed on the same day. In each plot, we recorded all plant species and estimated their cover. We identified plants to the species level, except for the genera *Crataegus, Epilobium, Rosa* and *Taraxacum*, which we recorded on genus level. For shrubs and trees, we estimated cover separately for the ground, shrub and tree layers.

To investigate the plant diversity, we used four different metrics: plant species richness (alpha diversity), the moisture index of the respective plants and dissimilarity on the section and landscape level (beta diversity). To assess the plant diversity on the section-level, we averaged the cover of all species across all plots within each section to obtain one compound relevé per section. We calculated the species richness as the number of distinct species per section. We quantified the moisture preference of plant communities in each section. We calculated a community-weighted plant moisture index using harmonized Ellenberg indicator values (Lubomír et al., 2022, hereafter “plant moisture index”). The preferences range from “drought tolerant” (F = 1) to “often submerged” (F = 11).

As measures of beta diversity, we characterized heterogeneity of the plant species composition on the section and landscape scale. At the section scale, we calculated Bray-Curtis dissimilarities based on the 4 m × 4 m vegetation plots. For each section, we calculated the dissimilarity of each individual plot to all other plots within the same section and averaged these values across all plots within the given section. We called it “within section dissimilarity”, and it quantifies how different the plant communities among plots within one section are from each other and this corresponds to within-site beta diversity. At the landscape scale, we calculated dissimilarities between the compound vegetation relevé of each section and the compound relevés of all other sections with the same restoration status (restored/control). We averaged these dissimilarities for a given section and called it “between section dissimilarity”. This quantifies how different the sections of one restoration status are from each other at the landscape level, and this corresponds to between-site beta diversity.

### 2.5. Statistical analysis

We processed and analyzed the data with R 4.4.1 (R Core Team, 2026). We evaluated assumptions of all models through diagnostic plots of residuals (Zuur & Ieno, 2016). For beta mixed-effect models we assessed the model fit with simulation-based residual diagnostics (R-package DHARMa, Hartig et al., 2024).

#### 2.5.1. Restoration effect on the response variables

To address the first hypothesis, the difference between restored and control sections, we tested if restoration affected stream structural complexity, habitat and plant diversity. We focused on seven ecological responses: stream structural complexity, habitat richness, proportion of key-floodplain habitats, plant species richness, plant moisture index, within section dissimilarity and between section dissimilarity. To test the effect of restoration on these response variables, we constructed a linear mixed-effect model for all variables except for the proportion of key floodplain habitats (*lmer* function, R-package *lme4*; Bates et al., 2015). We included the restoration status (restored vs. control) and the scaled section area as explanatory variables to quantify restoration effects while controlling for differences in section area. We included the pair identity number, which is shared between the restored and the corresponding control section as a random effect to account for their non-independence. As the proportion of key floodplain habitats is on a bounded scale between 0 and 1 and included true boundary values (0: n = 2; 1: n = 16), the assumption of normality in residuals would be violated for a gaussian distribution. We therefore fitted a beta mixed-effects model with logit-link (*glmmTMB* function, R-package *glmmTMB*, Brooks et al., 2017) and included section area and restoration status as explanatory variables and pair identity number as random effect (Douma & Weedon, 2019). Because beta distribution does not allow exact 0 or 1 values, we adjusted values slightly toward the interior while preserving relative ordering of values using the Smithson and Verkuilen transformation (Smithson & Verkuilen, 2006).

#### 2.5.2. Plant community composition

To test the second hypothesis, whether plant community composition differs between restored and control sections, we used Nonmetric Multidimensional Scaling (NMDS) based on the Bray-Curtis dissimilarity matrix of the species. We addressed zero-inflation by using Hellinger transformation and selected the number of dimensions and the number of random starts (trymax) to achieve stress < 0.2 with the lowest number of axes (Dexter et al., 2018). We tested if the community composition differed between restored and control sections using PERMANOVA (*adonis2* function, R-package *vegan*, Oksanen et al., 2025). We used the Bray-Curtis dissimilarity-matrix as the response variable and included restoration status as the predictor. Here we also accounted for the paired structure and used the pair identity as block factor. We assessed statistical significance with 199 permutations. We evaluated assumptions using the *betadisper* function of the R-package *vegan*. To identify species that are associated with restored or control sections, we performed an indicator species analysis (999 permutations, *multipatt* function, R-package *indicspecies*, Cáceres & Legendre, 2009). We kept species significantly associated (p ≤ 0.05) with control or restored sections, and with a minimum indicator value of 0.5. The indicator value ranges from 0 to 1 and reflects both the specificity and fidelity of a species to a group, with a higher value indicating a stronger association.

#### 2.5.3. Mediators of the restoration effect

To address the third hypothesis, whether stream structural complexity and habitat richness mediate the effect of restoration on biodiversity, we used structural equation modelling (*psem* function, R-package *piecewiseSEM* package, Lefcheck, 2016). Structural equation modelling allows simultaneous evaluation of direct and indirect pathways, and thus, allows to model multiple interacting processes in complex systems (Grace, 2006). We included the restoration status as a numerical variable (0 = control, 1 = restored) to represent the difference between restored and control sections and to simplify interpretation of the resulting path diagram.

The structural equation model consisted of five linear mixed-effect models and one beta mixed effect model (for the proportion of key floodplain habitats). We modeled the first two component models with stream structural complexity and habitat richness as potential mediators and related each to restoration status and section area. For habitat richness as response, we used stream structural complexity as an explanatory variable to check if there is a mediation effect too. Then we created four more component models with the proportion of key-floodplain habitats, plant species richness, plant moisture index, and dissimilarity between sections as response variables, and related them to section area, structural complexity, habitat richness and restoration status as explanatory variables. We included pair identity number as a random effect in all component models. The full model specification can be found in Table S3.

We evaluated the structural equation model to examine whether the hypothesized structure fits the data. We evaluated model adequacy using tests of direct separation implemented in the *psem* function (Lefcheck, 2016), which tests whether variables that are not linked in the hypothesized model are nevertheless statistically related. The test of direct separation revealed a missing causal path within our hypothesized model. We therefore extended the model by including the proportion of key floodplain habitats as an additional predictor of the plant moisture index in the corresponding component model. We assessed the overall model fit with Fisher’s *C* statistics, which follows a chi-squared distribution, where a p-value above 0.05 indicates that the hypothesized causal structure is consistent with the data, thus containing all important links. We calculated conditional and marginal R^2^ values to assess the variance explained by fixed factors only and fixed and random factors combined. To get standardized path coefficients, we scaled all variables prior to the analysis, except the proportion of key floodplain habitats. Because this variable is modelled using beta regression with logit-link, standardized path coefficients cannot be calculated (variance is dependent on the mean and therefore not constant as in gaussian distributions). We therefore report the odds ratio for the resulting paths, which quantifies how the odds of a higher chance for key floodplain habitats differ between restored and control sections.

#### 2.5.4. Development of restored sections over time

To address the fourth hypothesis, the development of restored sections over time, we analyzed how the stream structural complexity as well as plant and habitat diversity changed with time since restoration. Here, we worked with data on restored sections only. Since the time since restoration of the restored sites included in this study varied between 2 and 25 years, we employed a space-for-time approach. We focused on response variables for which we detected significant effects of restoration: stream structural complexity, habitat richness, proportion of key-floodplain habitats, plant species richness, plant moisture index and dissimilarity between sections. We related each response variable in a linear model to the section area (to account for it) and time since restoration. For the proportion of key floodplain habitats, we again used a beta regression model. Visual inspections indicated a non-linear relationship between plant species richness and time since restoration, so we tested a quadratic relationship using raw polynomial terms while checking variance inflation factors for signs of multicollinearity between linear and quadratic terms of time since restoration. We also tested all other models for quadratic relationships but none of the models increased their fit significantly when including quadratic terms. To allow quadratic terms, we centered the time since restoration prior to the analysis.

Sampling sites over a geographic range are often not spatially independent hence violating a core model assumption. This was not problematic when we compared restored and control sites, because their pairs were always located close to each other. However, when we tested the effect of time since restoration, we worked with the restored sections only.

Therefore, we tested all models that investigated the development of restored sites over time for residual spatial autocorrelation using Moran’s *I* plots to make sure that the assumptions of the models are met (Legendre & Legendre, 2012). We found significant spatial autocorrelation for the models with stream structural complexity (*I* = 0.42, p = 0.001) and plant moisture index (*I = 0.17,* p *= 0.02*) as response variables and addressed it by using spatial error models (Fig. S2). Spatial error models account for spatial autocorrelation by modeling the dependence of residuals among nearby observations in the error term of the model.

## 3. Results

### 3.1. Effect of restoration on the response variables

We recorded a total of 81 different habitats. Of these, 69 appeared in restored sites with 24 exclusive habitats and 57 in control sites with 12 exclusive ones. We identified six key-floodplain habitats (Table S4). We recorded a total of 331 herbaceous and woody plant species. Of these, we recorded 295 in restored sections, with 72 exclusive species, and 259 in control sections, with 36 exclusive species.

In line with our first two hypotheses, restored sections had higher structural richness and habitat and plant diversity than control sections: stream structural complexity (restored sections had +62% higher stream structural complexity than control sections), habitat richness (+18%), proportion of key-floodplain habitats (+45%), plant species richness (+20%), plant moisture index (+16%) and between section dissimilarity (+5%, Fig. 2A, Table 1). The only exception was dissimilarity within sections which did not differ between restored and control sections (Table 1).

**Figure 2:**
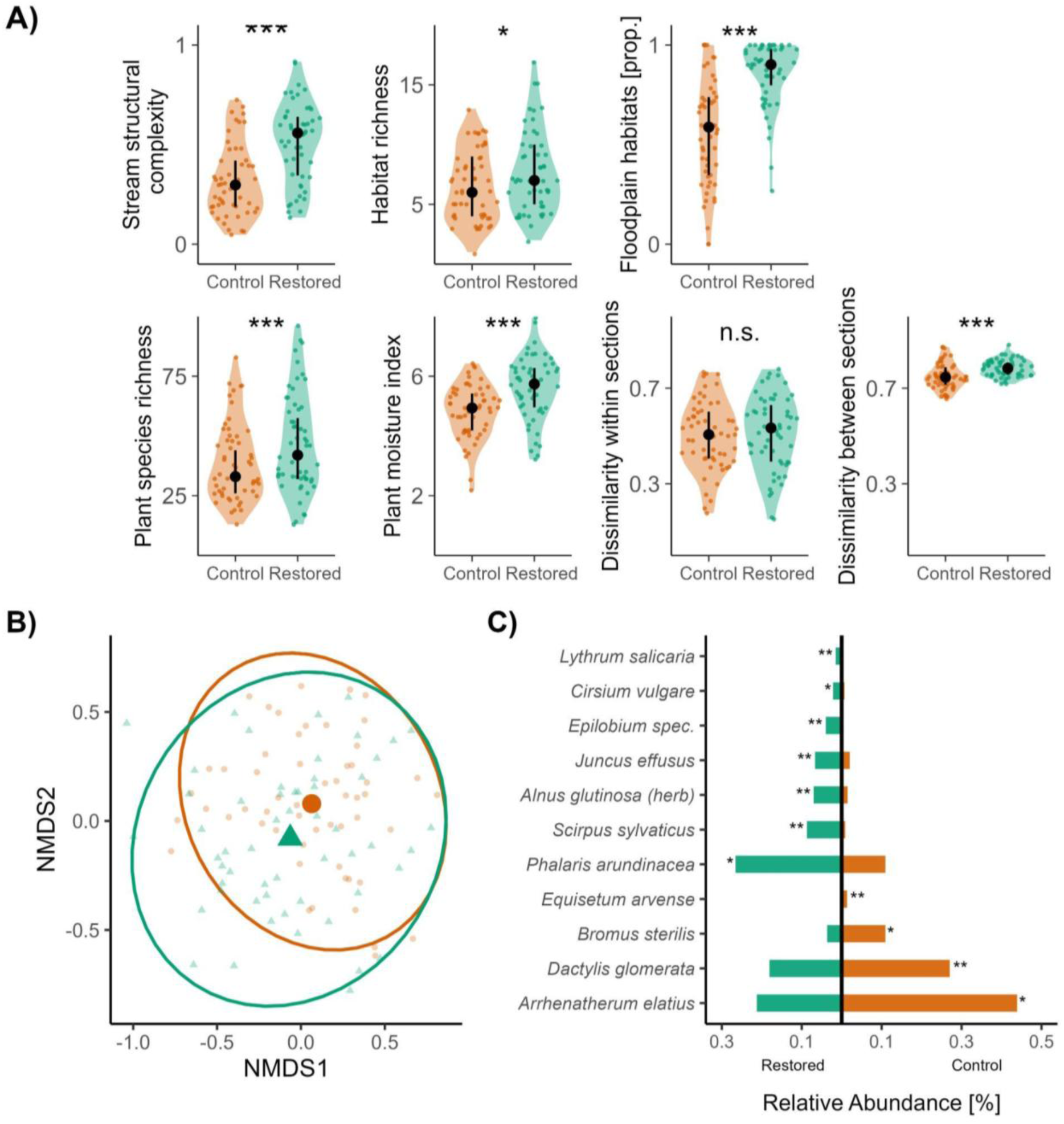
Comparison between restored and control sections. **A)** Stream structural complexity, habitat and plant diversity metrics. Points represent values for individual sections. Black dot is the median and black bars represent interquartile ranges (see Table 1 for model results). **B)** Composition of plant communities. Nonmetric multi-dimensional scaling (NMDS) with three-axis solution and stress = 0.169. Each point represents a section. See supplementary material (Fig. S1) for composition analysis including all three NMDS axes. **C)** Indicator species for restored and control sections with indicator values above 0.5. The bar represents relative abundance of species with significant associations in terms of specificity and fidelity to either restored or control sections. Asterisks indicate significant associations with given restoration status (*p < 0.05, **p < 0.01, ***p < 0.001).

**Table 1:**
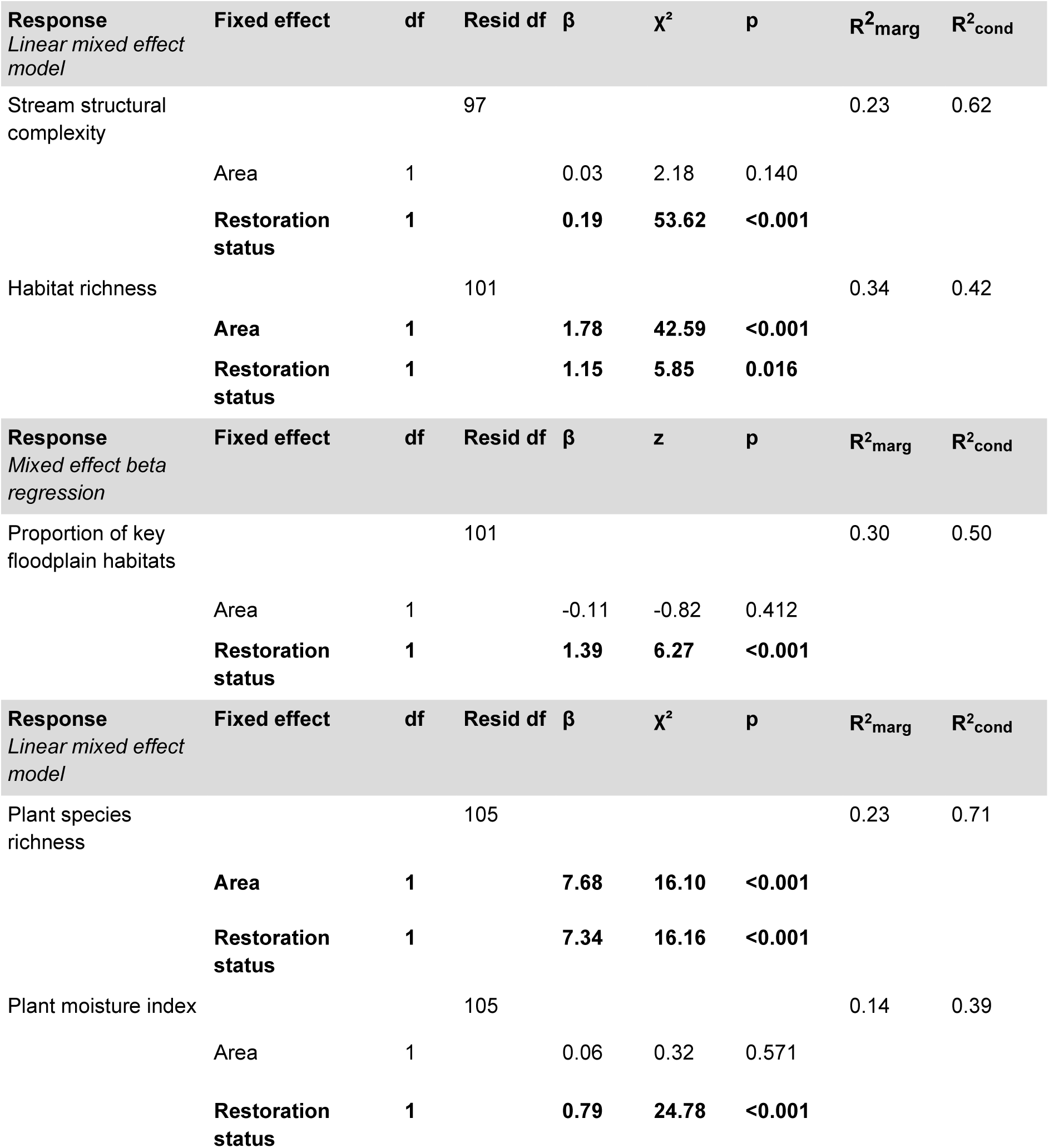
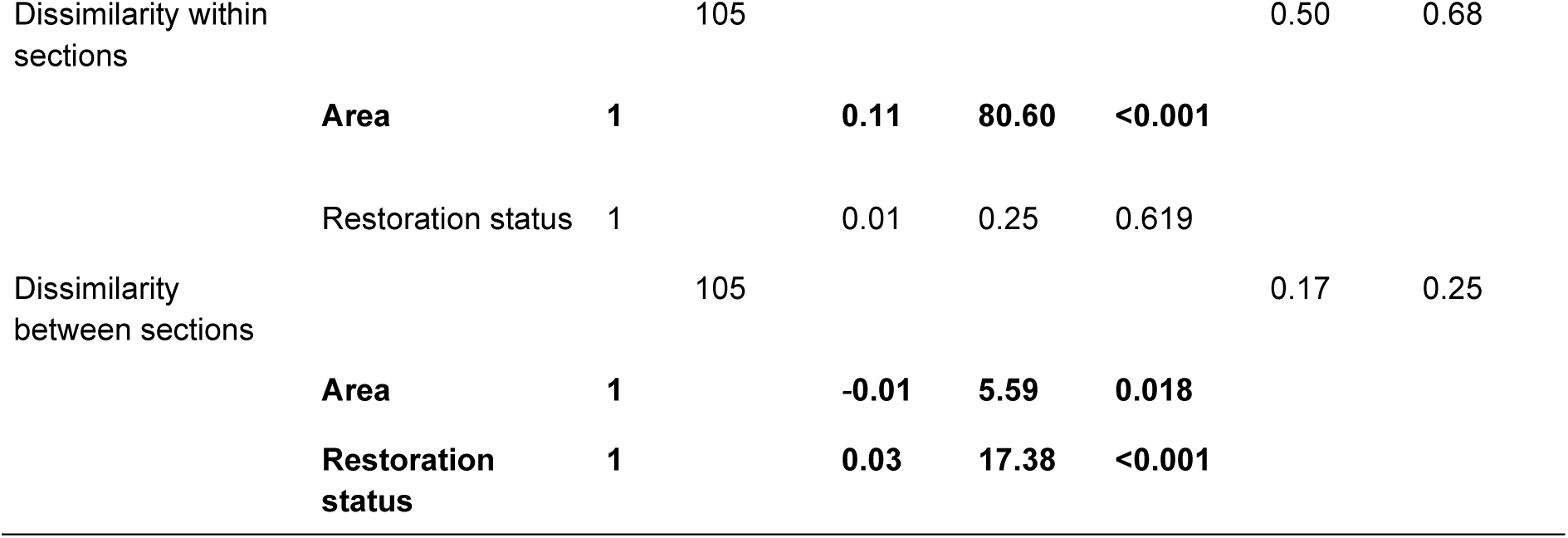
The effects of section area and restoration status (control vs. restored) on stream structural complexity, habitat richness, proportion of key-floodplain habitats, plant species richness, plant moisture index, dissimilarity within sections and dissimilarity between sections. We generated model output with linear mixed-effect models (pair identity number as random effect) except for the proportion of key floodplain habitats, where we used mixed effect beta regression. The respective model type is given in each header, and we report test statistics as corresponds to each model type. We obtained p-values for the fixed effects with Type II Analysis of Variance. Significant effects (p < 0.05) are highlighted in bold.

### 3.2. Plant community composition

The differences in plant diversity were only partially reflected by a shift in plant community composition. Nonmetric Multidimensional Scaling (three-axis solution with stress = 0.169, trymax = 200) visually showed no clear shifts in plant community composition between control and restored sites (Fig. 2B). PERMANOVA analysis indicated a significant differentiation of plant community composition by restoration status (F_1,108_ = 2.95, p = 0.005), but the explained variation was low (R^2^ = 0.03). When validating the assumptions of homoscedasticity, we found a difference in dispersion among our restoration statuses (F_1,108_ = 6.05, p = 0.016). The indicator species analysis identified 11 significant indicator species (p ≤ 0.05) that are associated with one of the restoration statuses (Fig. 2C). Seven species were associated with restored sections and four with control sections.

### 3.3. Mediators of the restoration effect

The third hypothesis predicted that the positive effects of restoration on biodiversity are mediated by increased stream structural complexity and habitat richness in restored sections. Restoration had indeed a strong direct positive effect on structural complexity (standardized path coefficient = 0.46) and habitat richness (0.18). Additionally, the model revealed the proportion of key floodplain habitats as an additional mediator (odds ratio: 2.60). These three variables acted as mediators of restoration effects (Fig. 3). Specifically, stream structural complexity positively affected the plant moisture index and the proportion of key floodplain habitats (standardized path coefficient = 0.37 and odds ratio: 1.58 respectively). This means that the increase of one standard deviation of structural complexity increased the plant moisture index by 0.37 standard deviations and increased the odds of a higher proportion of key floodplain habitats by 58%. Plant moisture index was mediated by two mediators: stream structural complexity and additionally the proportion of key floodplain habitats (standardized path coefficient = 0.31). Plant species richness was positively affected by habitat richness (standardized path coefficient = 0.34). Plant species richness had no effect on the plant moisture index. In contrast, the dissimilarity between sections was directly affected by restoration (standardized path coefficient = 0.31), but we did not measure a mediating effect. The structural equation model explained a substantial proportion of the variation in our data (Fig. 3) and indicated that all important paths are included (Fisher’s *C* = 6.968, df = 10, p = 0.728). In total, the component models explained 91% of the variation in key-floodplain habitats, 71% of the variation in plant species richness, 62% in structural complexity, 47% in plant moisture index, 41% in habitat richness and 27% in dissimilarity between sections.

**Figure 3:**
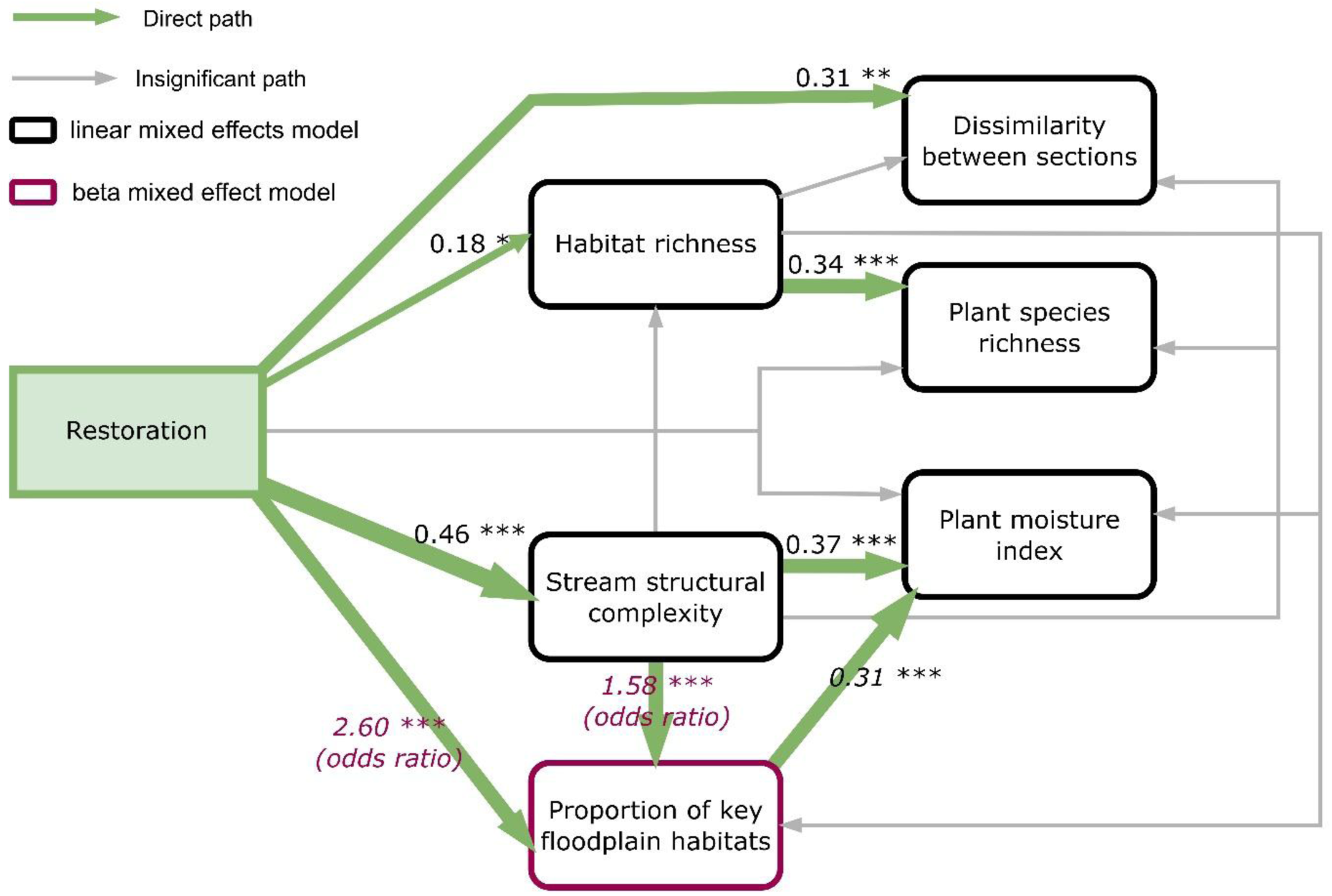
Path diagram of the structural equation model testing the direct and indirect effects of restoration on the response variables. Black values on the arrows are standardized path coefficients and represent the expected change in standard deviations in the response variable per one standard deviation change in the predictor while holding all other predictors constant. We used beta regression with logit-link for modelling the proportion of key floodplain habitats, purple values are therefore the change in odds ratio because standardized path coefficients are incalculable with logit-link. The change in odds ratio quantifies how the odds of a higher chance for key floodplain habitats differ between restored and control sections. The width of the arrows is proportional to the strength of the path coefficient or the change in odds ratio. All component models are corrected for the area of sections.

### 3.4. Development of restored sites over time

The fourth hypothesis predicted that habitat and plant diversity increases with time since restoration. Species richness in restored sections followed a U-shaped relationship with time since restoration, whereby it decreased 15 years followed by an increase thereafter (Table 2, Fig. 4). The proportion of key-floodplain habitats increased linearly with time since restoration, while habitat richness and dissimilarity between restored sections declined with time (Table 2, Fig. 4). Stream structural complexity and plant moisture index did not change with time since restoration (Fig. 4).

**Figure 4:**
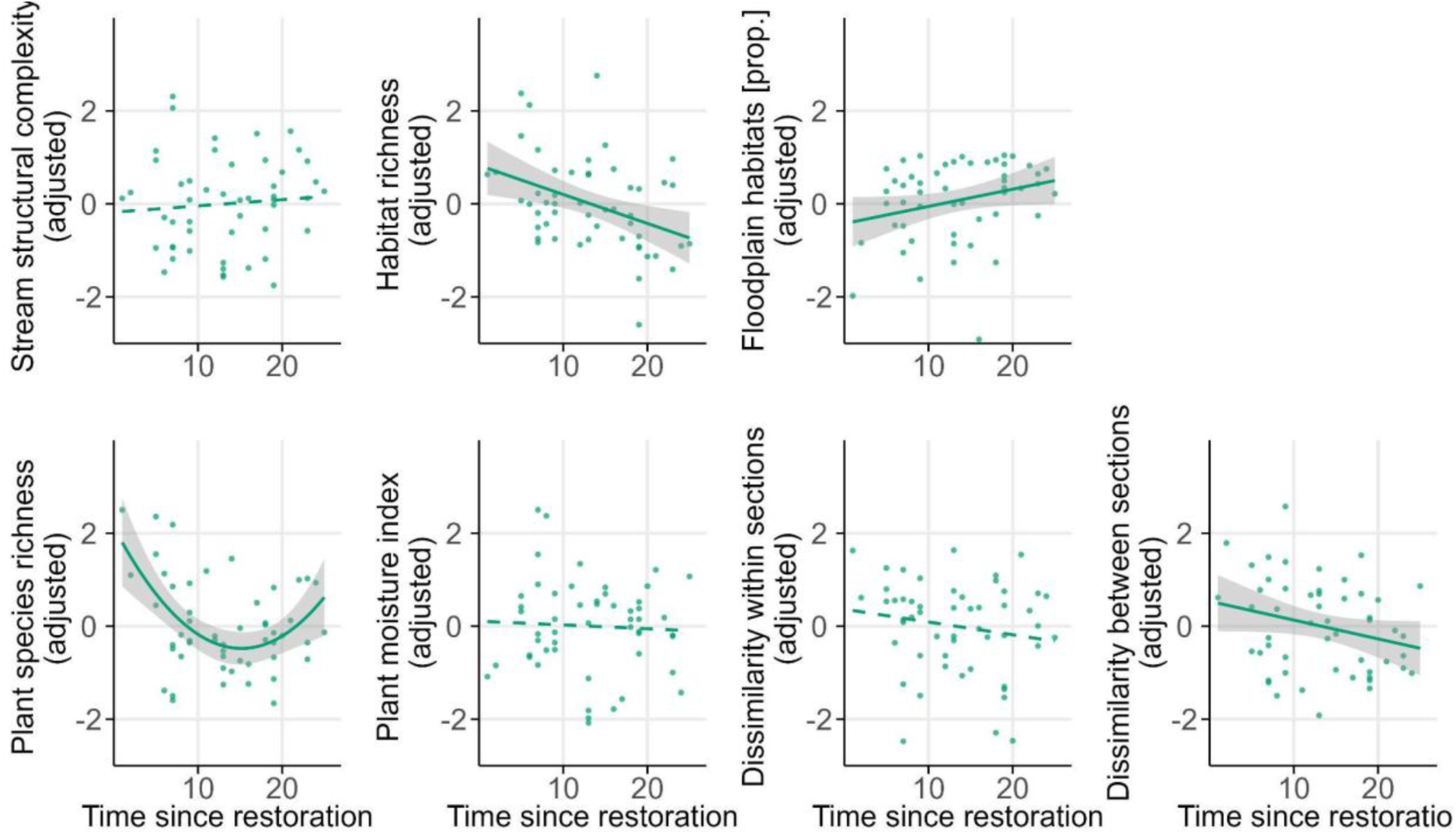
The effect of time since restoration on stream structural complexity and plant and habitat diversity and characteristics. Points represent values in individual restored sections. Solid lines and shaded confidence intervals represent significant relationships at p < 0.05; dashed lines indicate non-significant relationships. All response variables are adjusted for the area of given restored section. See Table 2 for model results.

**Table 2:**
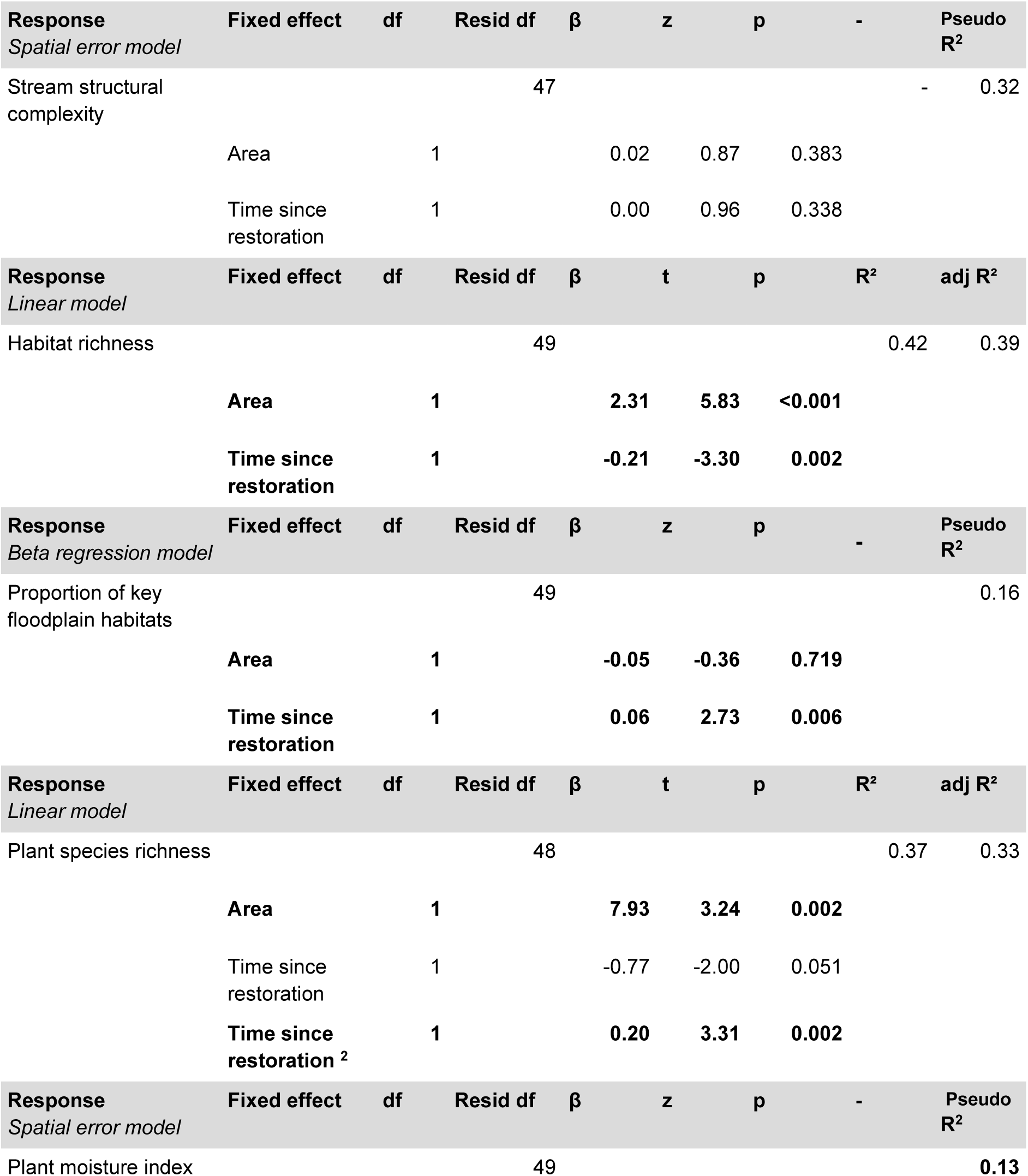
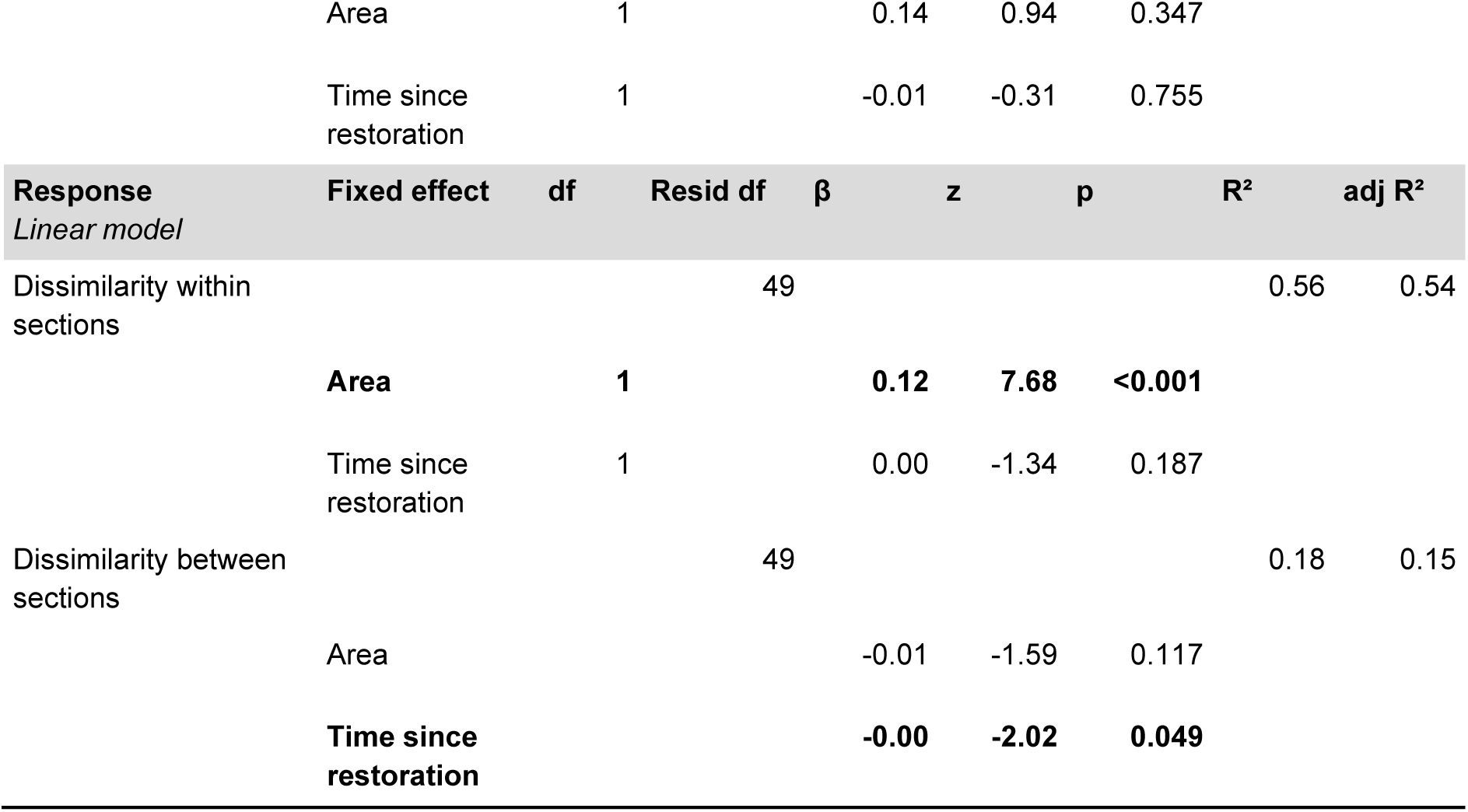
The effect of time since restoration on stream structural complexity, habitat richness, proportion of key-floodplain habitats, plant species richness, plant moisture index, dissimilarity within sections and dissimilarity between sections. These models include only restored sections. The respective model type is given in each header, and we report test statistics as corresponds to each model type. Significant effects (p < 0.05) are highlighted in bold.

| **Response**<br>*Spatial error model* | **Fixed effect** | **df** | **Resid df** | **$\beta$** | **z** | **p** | **-** | **Pseudo R<sup>2</sup>** |
| --- | --- | --- | --- | --- | --- | --- | --- | --- |
| Stream structural complexity |  |  | 47 |  |  |  | - | 0.32 |
|  | Area | 1 |  | 0.02 | 0.87 | 0.383 |  |  |
|  | Time since restoration | 1 |  | 0.00 | 0.96 | 0.338 |  |  |
| **Response**<br>*Linear model* | **Fixed effect** | **df** | **Resid df** | **$\beta$** | **t** | **p** | **R<sup>2</sup>** | **adj R<sup>2</sup>** |
| Habitat richness |  |  | 49 |  |  |  | 0.42 | 0.39 |
|  | **Area** | **1** |  | **2.31** | **5.83** | **<0.001** |  |  |
|  | **Time since restoration** | **1** |  | **-0.21** | **-3.30** | **0.002** |  |  |
| **Response**<br>*Beta regression model* | **Fixed effect** | **df** | **Resid df** | **$\beta$** | **z** | **p** | **-** | **Pseudo R<sup>2</sup>** |
| Proportion of key floodplain habitats |  |  | 49 |  |  |  |  | 0.16 |
|  | **Area** | **1** |  | **-0.05** | **-0.36** | **0.719** |  |  |
|  | **Time since restoration** | **1** |  | **0.06** | **2.73** | **0.006** |  |  |
| **Response**<br>*Linear model* | **Fixed effect** | **df** | **Resid df** | **$\beta$** | **t** | **p** | **R<sup>2</sup>** | **adj R<sup>2</sup>** |
| Plant species richness |  |  | 48 |  |  |  | 0.37 | 0.33 |
|  | **Area** | **1** |  | **7.93** | **3.24** | **0.002** |  |  |
|  | Time since restoration | 1 |  | -0.77 | -2.00 | 0.051 |  |  |
|  | **Time since restoration<sup>2</sup>** | **1** |  | **0.20** | **3.31** | **0.002** |  |  |
| **Response**<br>*Spatial error model* | **Fixed effect** | **df** | **Resid df** | **$\beta$** | **z** | **p** | **-** | **Pseudo R<sup>2</sup>** |
| Plant moisture index |  |  | 49 |  |  |  |  | 0.13 |

|  |  |  |  |  |
| --- | --- | --- | --- | --- |
| Area | 1 | 0.14 | 0.94 | 0.347 |
| Time since restoration | 1 | -0.01 | -0.31 | 0.755 |

| Response | Fixed effect | df | Resid df | $\beta$ | z | p | R<sup>2</sup> | adj R<sup>2</sup> |
| --- | --- | --- | --- | --- | --- | --- | --- | --- |
| *Linear model* |  |  |  |  |  |  |  |  |
| Dissimilarity within sections |  |  | 49 |  |  |  | 0.56 | 0.54 |
|  | **Area** | **1** |  | **0.12** | **7.68** | **<0.001** |  |  |
|  | Time since restoration | 1 |  | 0.00 | -1.34 | 0.187 |  |  |
| Dissimilarity between sections |  |  | 49 |  |  |  | 0.18 | 0.15 |
|  | Area |  |  | -0.01 | -1.59 | 0.117 |  |  |
|  | **Time since restoration** |  |  | **-0.00** | **-2.02** | **0.049** |  |  |

Source: Hessian Agency for Nature Conservation, Environment and Geology (HLNUG), 2022, license: CC BY 4.0.

## 4. Discussion

### 4.1. Small stream restoration increases stream structural complexity, habitat and plant diversity on a local and landscape scale

Restoration of small streams is common, but its effect on riparian biodiversity has been poorly documented since most studies focus on large rivers. Our study provides rare insights into the effects of restoration on small stream habitats and associated terrestrial biota. We show across 55 small streams in intensively used agricultural landscapes, that restoration consistently increased stream structural complexity and habitat richness. These physical changes have led to a higher cover of typical floodplain habitats and a positive response of riparian vegetation, including higher species richness and landscape heterogeneity. Our results highlight that the restoration of small streams can have a positive impact on habitat features and can support plant diversity in the agricultural landscape.

Our results show that small stream restoration increases stream structural complexity, habitat and plant diversity, which is in line with our first hypothesis. Restored sections had more habitats, with a higher proportion of key floodplain habitats and more plants with higher plant moisture index values. This indicates a shift towards more riparian conditions and improved lateral connectivity. Similar increases in floodplain habitat extent and wetland characteristics after restoration have also been reported for large to medium sized rivers (Januschke et al., 2011 and Lorenz et al., 2018). At the landscape level, restoration increased plant species richness (alpha diversity) and heterogeneity on the landscape scale (beta diversity, dissimilarity between sections). This suggests that restoration did not lead to uniform outcomes but instead produced a mosaic of site conditions across the landscape, a pattern which was already documented for stream restoration (see also Fraaije et al., 2019). Such spatial heterogeneity likely reflects differences in applied restoration measures, the local geomorphology, hydrological connectivity and the surrounding land use (Palmer et al., 2010; Suding, 2011).

### 4.2. Restoration increases variability in species composition rather than community turnover

Restoration only weakly influenced community composition. Despite a statistical difference in community composition, small effect sizes, large compositional overlap (NMDS ordination), and significant differences in dispersion of the within-group variability indicate that restoration mainly increased the variability in species composition rather than changing the whole communities per se. Thus, restored sites still draw from the same regional species pool but assemble differently depending on the local conditions. Indeed, other studies reported similarly weak compositional changes after restoration of riparian vegetation in rivers and streams (Kuglerová et al., 2017; Wenskus et al., 2025). Together, these results show that restoration increases biodiversity primarily by promoting spatial heterogeneity rather than by replacing the entire plant communities.

### 4.3. Structural complexity, habitat diversity and key floodplain habitats mediate the diversity responses to restoration

The effect of restoration on biodiversity was largely mediated through the increased stream structural complexity and increased habitat richness, parameters that are often targeted by restoration measures. This is in line with the habitat heterogeneity hypothesis, which predicts that biodiversity increases with increasing habitat heterogeneity (Stein et al., 2014; Tews et al., 2004). Restoration created additional habitats that expanded the available niche space. Similar indirect effects have been reported in large rivers where plant species richness primarily responded to greater habitat availability (Januschke et al., 2014; Wintle & Kirkpatrick, 2007). Other studies from small streams show that riparian vegetation richness strongly depends on habitat richness (Corenblit et al., 2007; Ye et al., 2024).

Restoration also increased the plant moisture index through changes in hydrological conditions through two linked pathways. On the one hand, a higher stream complexity directly promoted higher moisture values of the riparian vegetation, likely through restored lateral and vertical connectivity and increased variation in water tables and flooding regimes (Wenskus et al., 2025). On the other hand, restoration increased the proportion of key floodplain habitats both directly and indirectly through structural complexity. As floodplain habitats themselves host more moisture adapted species, they further increase the plant moisture index. This hydrological shift is also reflected by the indicator species of restored sites that are associated with wetter habitats, such as *Alnus glutinosa* and *Scirpus sylvaticus*. Similar increases of plant moisture indexes have been reported for rivers (Modrak et al., 2017) and for small streams (Driscoll et al., 2025). In contrast, several studies reported no differences in plant moisture indexes between restored and control sites for rivers and streams, probably because hydrological reconnection remained weak (Fraaije et al., 2019; Januschke et al., 2011). Our results show that vegetation responds to structural enhancement when restoration improves stream structure and increases water availability in the floodplain.

### 4.4. Successional processes continue after physical habitat conditions have stabilized

The responses of habitat and plant diversity to time since restoration varied among the different metrics. We hypothesized that plant and habitat diversity would increase with time since restoration as hydro morphological dynamics increasingly shape the floodplain landscape. Our results only partially supported this expectation. Plant species richness followed a U-shaped pattern with the highest richness in young restorations, a decline during intermediate stages, and a subsequent increase in older sections. Early peaks likely reflect the colonization by pioneer and disturbance adapted species, while the later increase suggests gradual establishment of later-successional species such as the tree *Alnus glutinosa* (Bauer et al., 2018; Noon, 1996). This is also indicated by the increase of key floodplain habitats in older sections. In contrast to our hypothesis, habitat richness and dissimilarity between sections declined linearly with time since restoration, indicating progressive habitat stabilization and the loss of early successional habitat types, which reduces compositional differences among sections. Importantly, this decline does not indicate ecological degradation. Instead, it reflects a directional shift toward a lower number of characteristic and well-developed floodplain habitats, such as softwood riparian forests. Stream structural complexity and the plant moisture index remained stable throughout the 2 to 25 years since restoration, indicating that physical and hydrological conditions stabilized shortly after restoration and stayed stable over time. This contrasts with our expectation that stream structural complexity will increase over time due to reestablished floodplain dynamics. Possibly, the small streams have insufficient discharge to allow post-restoration channel dynamics. Additionally, many restored streams are still confined by surrounding land use and persistent catchment-scale pressures, particularly agricultural land use and nutrient inputs (Amoros & Bornette, 2002; Muhar et al., 2016). These results suggest that vegetation continues to reorganize through successional processes even after physical channel conditions have stabilized.

### 4.5. Restoration outcomes appear transferable across river sizes, but temporal patterns differ

We have found largely the same habitat and plant diversity responses to restoration in small streams as reported for large rivers. The only difference was their development over time. In large rivers, species richness and habitat richness often show weak or no relationships with time since restoration (Modrak et al., 2017; Pilotto et al., 2019). In contrast, the few existing studies on smaller streams have reported relationships between vegetation responses and time since restoration (Hasselquist et al., 2015; Lennox et al., 2011), which is consistent with findings from our study. Small streams experience more frequent but shorter flooding events and operate under tighter spatial constraints, which can lead to faster ecological responses once structural conditions are restored (Dietrich et al., 2014). In our study, riparian succession was dominated by *Alnus glutinosa*, reflecting a typical development pathway of small stream floodplain forests. Our results therefore suggest that vegetation dynamics in small streams are driven by rather rapid successional processes than by long-term geomorphic alterations which might appear in rivers and take longer time spans.

Consequently, the positive responses of habitat and plant diversity appear transferable from rivers to streams, whereas the temporal patterns are not. Overall, our results suggest that stream restoration can successfully re-establish characteristic riparian habitats and support associated plant diversity, highlighting its importance for the recovery of ecosystems that have largely disappeared from agricultural landscapes.

## 5. Implications for practitioners

Restoration increases structural complexity, habitat diversity and reconnects the floodplains with their streams, which has cascading positive effects on plant diversity and riparian moisture conditions. Our study therefore shows that small stream restoration can be effective in increasing biodiversity around streams in agricultural landscapes. Restoration even increases plant beta diversity on the landscape scale and thus effectively reduces biotic homogenization. Our results suggest that restoration enhances biodiversity primarily by increasing stream structural complexity and habitat richness. Wherever possible, restoration practitioners should therefore focus on substantial improvement of the stream structure and creation of diverse habitats. This is particularly important in small streams, where the restored structures persist through time because the water discharge is insufficient to remodel the channel following the restoration.

## Supporting information

Lerbs_et_al_supplementary_material

## Acknowledgement

This project was funded by Lore-Steubing-Institute as part of the HLNUG (Hessisches Landesamt Naturschutz, Umwelt und Geologie).

## Authors contributions

AB and SP conceived the idea with support from SL and NF; AB and LL designed the study; LL, AD, LG, AS and JL collected the data; LL curated the data; LL and FW conducted the formal analysis; LL wrote the first draft with support from AB, all authors edited the manuscript and approved the final version.

## References

1. Amoros, C., & Bornette, G. (2002). Connectivity and biocomplexity in waterbodies of riverine floodplains. Freshwater Biology, 47(4), 761–776. 10.1046/j.1365-2427.2002.00905.x

2. Bates, D., Mächler, M., Bolker, B., & Walker, S. (2015). Fitting Linear Mixed-Effects Models Using lme4. Journal of Statistical Software, 67, 1–48. 10.18637/jss.v067.i01

3. Bauer, M., Harzer, R., Strobl, K., & Kollmann, J. (2018). Resilience of riparian vegetation after restoration measures on River Inn. River Research and Applications, 34(5), 451–460. 10.1002/rra.3255

4. Baumane, M., Baastrup-Spohr, L., Sand-Jensen, K., Goldberg, I., Thorø Martinsen, K., & Bruun, H. H. (2024). Nutrients, isolation and lack of grazing limit plant diversity in restored wetlands. Journal of Applied Ecology, n/a(n/a). 10.1111/1365-2664.14824

5. Bernhardt, E. S., Palmer, M. A., Allan, J. D., Alexander, G., Barnas, K., Brooks, S., Carr, J., Clayton, S., Dahm, C., Follstad-Shah, J., Galat, D., Gloss, S., Goodwin, P., Hart, D., Hassett, B., Jenkinson, R., Katz, S., Kondolf, G. M., Lake, P. S., … Sudduth, E. (2005). Synthesizing U.S. River Restoration Efforts. Science, 308(5722), 636–637. 10.1126/science.1109769

6. Brooks, M. E., Kristensen, K., Benthem, K. J. van, Magnusson, A., Berg, C. W., Nielsen, A., Skaug, H. J., Mächler, M., & Bolker, B. M. (2017). glmmTMB Balances Speed and Flexibility Among Packages for Zero-inflated Generalized Linear Mixed Modeling. The R Journal, 9(2), 378–400. 10.32614/RJ-2017-066

7. Cáceres, M. D., & Legendre, P. (2009). Associations between species and groups of sites: Indices and statistical inference. Ecology, 90(12), 3566–3574. 10.1890/08-1823.1

8. Corenblit, D., Tabacchi, E., Steiger, J., & Gurnell, A. M. (2007). Reciprocal interactions and adjustments between fluvial landforms and vegetation dynamics in river corridors: A review of complementary approaches. Earth-Science Reviews, 84(1), 56–86. 10.1016/j.earscirev.2007.05.004

9. Dietrich, A. L., Lind, L., Nilsson, C., & Jansson, R. (2014). The Use of Phytometers for Evaluating Restoration Effects on Riparian Soil Fertility. Journal of Environmental Quality, 43(6), 1916–1925. 10.2134/jeq2014.05.0197

10. Douma, J. C., & Weedon, J. T. (2019). Analysing continuous proportions in ecology and evolution: A practical introduction to beta and Dirichlet regression. Methods in Ecology and Evolution, 10(9), 1412–1430. 10.1111/2041-210X.13234

11. Driscoll, K. P., Martinez, L. F., & Turner, T. F. (2025). Stream restoration effectively alters functional diversity and composition of riparian plant communities in the southern Rocky Mountains, U.S.A. Restoration Ecology, 33(5), e70053. 10.1111/rec.70053

12. Dudgeon, D., Arthington, A. H., Gessner, M. O., Kawabata, Z.-I., Knowler, D. J., Lévêque, C., Naiman, R. J., Prieur-Richard, A.-H., Soto, D., Stiassny, M. L. J., & Sullivan, C. A. (2006). Freshwater biodiversity: Importance, threats, status and conservation challenges. Biological Reviews, 81(2), 163–182. 10.1017/S1464793105006950

13. Fraaije, R. G. A., Poupin, C., Verhoeven, J. T. A., & Soons, M. B. (2019). Functional responses of aquatic and riparian vegetation to hydrogeomorphic restoration of channelized lowland streams and their valleys. Journal of Applied Ecology, 56(4), 1007–1018. 10.1111/1365-2664.13326

14. Google LLC. (2024). Google Earth (Version 7.3.6.9796) [Software]. https://earth.google.com/earth/

15. Grace, J. B. (2006). Structural Equation Modeling and Natural Systems. Cambridge University Press. 10.1017/CBO9780511617799

16. Hartig, F., Lohse, L., & leite, M. de S. (2024). DHARMa: Residual Diagnostics for Hierarchical (Multi-Level / Mixed) Regression Models (Version 0.4.7) [Software]. https://cran.r-project.org/web/packages/DHARMa/index.html

17. Hasselquist, E. M., Nilsson, C., Hjältén, J., Jørgensen, D., Lind, L., & Polvi, L. E. (2015). Time for recovery of riparian plants in restored northern Swedish streams: A chronosequence study. Ecological Applications, 25(5), 1373–1389. 10.1890/14-1102.1

18. Hering, D., Aroviita, J., Baattrup-Pedersen, A., Brabec, K., Buijse, T., Ecke, F., Friberg, N., Gielczewski, M., Januschke, K., Köhler, J., Kupilas, B., Lorenz, A. W., Muhar, S., Paillex, A., Poppe, M., Schmidt, T., Schmutz, S., Vermaat, J., Verdonschot, P. F. M., … Kail, J. (2015). Contrasting the roles of section length and instream habitat enhancement for river restoration success: A field study of 20 European restoration projects. Journal of Applied Ecology, 52(6), 1518–1527. 10.1111/1365-2664.12531

19. Hessisches Wassergesetz. (2010). https://www.rv.hessenrecht.hessen.de/bshe/document/jlr-WasGHE2010pP2 (accessed: 07.01.2025)

20. Januschke, K., Brunzel, S., Haase, P., & Hering, D. (2011). Effects of stream restorations on riparian mesohabitats, vegetation and carabid beetles. Biodiversity and Conservation, 20(13), 3147–3164. 10.1007/s10531-011-0119-8

21. Januschke, K., Hering, D., Scholz, M., Ehlert, T., Rumm, A., & Stammel, B. (2024). Biozönotische Erfolgskontrolle von Renaturierungsmaßnahmen an Gewässerufern und in Auen. Wasser und Abfall, 26(6), 36–41. 10.1007/s35152-024-1865-8

22. Januschke, K., Jähnig, S. C., Lorenz, A. W., & Hering, D. (2014). Mountain river restoration measures and their success(ion): Effects on river morphology, local species pool, and functional composition of three organism groups. Ecological Indicators, 38, 243–255. 10.1016/j.ecolind.2013.10.031

23. Kollmann, J., Kirmer, A., Tischew, S., Hölzel, N., & Kiehl, K. (2019). Renaturierungsökologie. Springer-Verlag.

24. Kristensen, P., & Globevnik, L. (2014). European small water bodies. Biology and Environment: Proceedings of the Royal Irish Academy, 114B(3), 281–287. 10.3318/BIOE.2014.13

25. Kuglerová, L., Botková, K., & Jansson, R. (2017). Responses of riparian plants to habitat changes following restoration of channelized streams. Ecohydrology, 10(1), e1798. 10.1002/eco.1798

26. LAWA (1999). Gewässerstrukturgütekartierung in der Bundesrepublik Deutschland. https://www.lawa.de/documents/gewaesserstrukturguetekartierung_verfahren_kleine_mittelgrosse_fliessgewaesser_1552305499.pdf (accessed: 17.11.2023)

27. Lefcheck, J. S. (2016). piecewiseSEM: Piecewise structural equation modelling in r for ecology, evolution, and systematics. Methods in Ecology and Evolution, 7(5), 573–579. 10.1111/2041-210X.12512

28. Legendre, P., & Legendre, L. (2012). Numerical Ecology. Elsevier Science.

29. Lennox, M. S., Lewis, D. J., Jackson, R. D., Harper, J., Larson, S., & Tate, K. W. (2011). Development of Vegetation and Aquatic Habitat in Restored Riparian Sites of California’s North Coast Rangelands. Restoration Ecology, 19(2), 225–233. 10.1111/j.1526-100X.2009.00558.x

30. Lepori, F., Palm, D., Brännäs, E., & Malmqvist, B. (2005). Does Restoration of Structural Heterogeneity in Streams Enhance Fish and Macroinvertebrate Diversity? Ecological Applications, 15(6), 2060–2071. 10.1890/04-1372

31. Leuschner, C., & Ellenberg, H. (2017). Ecology of Central European Forests: Vegetation Ecology of Central Europe, Volume I. Springer.

32. Lorenz, A. W., & Feld, C. K. (2013). Upstream river morphology and riparian land use overrule local restoration effects on ecological status assessment. Hydrobiologia, 704(1), 489–501. 10.1007/s10750-012-1326-3

33. Lorenz, A. W., Haase, P., Januschke, K., Sundermann, A., & Hering, D. (2018). Revisiting restored river reaches – Assessing change of aquatic and riparian communities after five years. Science of The Total Environment, 613–614, 1185–1195. 10.1016/j.scitotenv.2017.09.188

34. Lubomír, T., Axmanová, I., Dengler, J., Guarino, R., Jansen, F., Midolo, G., Nobis, M. P., Meerbeek, K. V., Aćić, S., Attorre, F., Bergmeier, E., Biurrun, I., Bonari, G., Bruelheide, H., Campos, J. A., Čarni, A., Chiarucci, A., Ćuk, M., Ćušterevska, R., … Chytrý, M. (2022). Ellenberg-type indicator values for European vascular plant species. 10.5281/zenodo.7427088

35. Meyer, J. L., Strayer, D. L., Wallace, J. B., Eggert, S. L., Helfman, G. S., & Leonard, N. E. (2007). The Contribution of Headwater Streams to Biodiversity in River Networks. JAWRA Journal of the American Water Resources Association, 43(1), 86–103. 10.1111/j.1752-1688.2007.00008.x

36. Modrak, P., Brunzel, S., & Lorenz, A. W. (2017). Riparian plant species preferences indicate diversification of site conditions after river restoration. Ecohydrology, 10(5), e1852. 10.1002/eco.1852

37. Morandi, B., Kail, J., Toedter, A., Wolter, C., & Piégay, H. (2017). Diverse Approaches to Implement and Monitor River Restoration: A Comparative Perspective in France and Germany. Environmental Management, 60(5), 931–946. 10.1007/s00267-017-0923-3

38. Muhar, S., Januschke, K., Kail, J., Poppe, M., Schmutz, S., Hering, D., & Buijse, A. D. (2016). Evaluating good-practice cases for river restoration across Europe: Context, methodological framework, selected results and recommendations. Hydrobiologia, 769(1), 3–19. 10.1007/s10750-016-2652-7

39. Naiman, R. J., & Décamps, H. (1997). The Ecology of Interfaces: Riparian Zones. Annual Review of Ecology, Evolution, and Systematics, 28(Volume 28, 1997), 621–658. 10.1146/annurev.ecolsys.28.1.621

40. Naiman, R. J., Décamps, H., & McClain, M. E. (2013). Riparian Landscapes. In Encyclopedia of Biodiversity (S. 461–468). Elsevier. 10.1016/B978-0-12-384719-5.00393-2

41. Nilsson, C., Polvi, L. E., Gardeström, J., Hasselquist, E. M., Lind, L., & Sarneel, J. M. (2015). Riparian and in-stream restoration of boreal streams and rivers: Success or failure? Ecohydrology, 8(5), 753–764. 10.1002/eco.1480

42. Noon, K. F. (1996). A model of created wetland primary succession. Landscape and Urban Planning, 34(2), 97–123. 10.1016/0169-2046(95)00209-X

43. Oksanen, J., Simpson, G. L., Blanchet, F. G., Kindt, R., Legendre, P., Minchin, P. R., O’Hara, R. B., Solymos, P., Stevens, M. H. H., Szoecs, E., Wagner, H., Barbour, M., Bedward, M., Bolker, B., Borcard, D., Borman, T., Carvalho, G., Chirico, M., Caceres, M. D., … Weedon, J. (2025). vegan: Community Ecology Package (Version 2.7-2) [Software]. https://cran.r-project.org/web/packages/vegan/index.html

44. Palmer, M. A., Menninger, H. L., & Bernhardt, E. (2010). River restoration, habitat heterogeneity and biodiversity: A failure of theory or practice? Freshwater Biology, 55(s1), 205–222. 10.1111/j.1365-2427.2009.02372.x

45. Pickett, S. T. A. (1989). Space-for-Time Substitution as an Alternative to Long-Term Studies. In G. E. Likens (Hrsg.), Long-Term Studies in Ecology: Approaches and Alternatives (S. 110–135). Springer. 10.1007/978-1-4615-7358-6_5

46. Pilotto, F., Tonkin, J. D., Januschke, K., Lorenz, A. W., Jourdan, J., Sundermann, A., Hering, D., Stoll, S., & Haase, P. (2019). Diverging response patterns of terrestrial and aquatic species to hydromorphological restoration. Conservation Biology, 33(1), 132–141. 10.1111/cobi.13176

47. QGIS Development Team. (2023). QGIS Geographic Information System (Version 3.28.8-Firenze) [Software]. QGIS Association. https://www.qgis.org

48. R Core Team. (2026). R: A Language and Environment for Statistical Computing [Software]. R Foundation for Statistical Computing. https://www.r-project.org/

49. Reid, A. J., Carlson, A. K., Creed, I. F., Eliason, E. J., Gell, P. A., Johnson, P. T. J., Kidd, K. A., MacCormack, T. J., Olden, J. D., Ormerod, S. J., Smol, J. P., Taylor, W. W., Tockner, K., Vermaire, J. C., Dudgeon, D., & Cooke, S. J. (2019). Emerging threats and persistent conservation challenges for freshwater biodiversity. Biological Reviews, 94(3), 849–873. 10.1111/brv.12480

50. Riis, T., Kelly-Quinn, M., Aguiar, F. C., Manolaki, P., Bruno, D., Bejarano, M. D., Clerici, N., Fernandes, M. R., Franco, J. C., Pettit, N., Portela, A. P., Tammeorg, O., Tammeorg, P., Rodríguez-González, P. M., & Dufour, S. (2020). Global Overview of Ecosystem Services Provided by Riparian Vegetation. BioScience, 70(6), 501–514. 10.1093/biosci/biaa041

51. Schubert, R., Klotz, S., & Hilbig, W. (2010). Bestimmungsbuch der Pflanzengesellschaften Deutschlands. https://link.springer.com/book/9783827425843

52. Smithson, M., & Verkuilen, J. (2006). A better lemon squeezer? Maximum-likelihood regression with beta-distributed dependent variables. Psychological Methods, 11(1), 54–71. 10.1037/1082-989X.11.1.54

53. Stein, A., Gerstner, K., & Kreft, H. (2014). Environmental heterogeneity as a universal driver of species richness across taxa, biomes and spatial scales. Ecology Letters, 17(7), 866–880. 10.1111/ele.12277

54. Suding, K. N. (2011). Toward an Era of Restoration in Ecology: Successes, Failures, and Opportunities Ahead. Annual Review of Ecology, Evolution, and Systematics, 42(1), 465–487. 10.1146/annurev-ecolsys-102710-145115

55. Tews, J., Brose, U., Grimm, V., Tielbörger, K., Wichmann, M. C., Schwager, M., & Jeltsch, F. (2004). Animal species diversity driven by habitat heterogeneity/diversity: The importance of keystone structures. Journal of Biogeography, 31(1), 79–92. 10.1046/j.0305-0270.2003.00994.x

56. Tockner, K., & Stanford, J. A. (2002). Riverine flood plains: Present state and future trends. Environmental Conservation, 29(3), 308–330. 10.1017/S037689290200022X

57. Vannote, R. L., Minshall, G. W., Cummins, K. W., Sedell, J. R., & Cushing, C. E. (1980). The River Continuum Concept. Canadian Journal of Fisheries and Aquatic Sciences, 37(1), 130–137. 10.1139/f80-017

58. Verdonschot, P. F. M., & Verdonschot, R. C. M. (2023). The role of stream restoration in enhancing ecosystem services. Hydrobiologia, 850(12), 2537–2562. 10.1007/s10750-022-04918-5

59. Ward, J. v., Tockner, K., & Schiemer, F. (1999). Biodiversity of floodplain river ecosystems: Ecotones and connectivity. Regulated Rivers: Research & Management, 15(1–3), 125–139. 10.1002/(SICI)1099-1646(199901/06)15:1/3%3C125::AID-RRR523%3E3.0.CO;2-E

60. Wenskus, F., Hecht, C., Hering, D., Januschke, K., Rieland, G., Rumm, A., Scholz, M., Weber, A., & Horchler, P. (2025). Effects of floodplain decoupling on taxonomic and functional diversity of terrestrial floodplain organisms. Ecological Indicators, 170, 113106. 10.1016/j.ecolind.2025.113106

61. Wintle, B. C., & Kirkpatrick, J. B. (2007). The response of riparian vegetation to flood-maintained habitat heterogeneity. Austral Ecology, 32(5), 592–599. 10.1111/j.1442-9993.2007.01753.x

62. Ye, M., Hu, H., Wu, P., Xie, Z., Hu, Y., & Lu, X. (2024). Ecological responses to hydrological connectivity in grassland riparian zones: Insights from vegetation and ground-dwelling arthropods. Science of The Total Environment, 922, 171196. 10.1016/j.scitotenv.2024.171196

63. Zerbe, S., & Wiegleb, G. (Hrsg.). (2009). Renaturierung von Ökosystemen in Mitteleuropa. Springer Berlin Heidelberg. 10.1007/978-3-662-48517-0

64. Zuur, A. F., & Ieno, E. N. (2016). A protocol for conducting and presenting results of regression-type analyses. Methods in Ecology and Evolution, 7(6), 636–645. 10.1111/2041-210X.12577

