## Supplementary material for "Small stream restoration increases habitat and plant diversity across scales": Lerbs_et_al_supplementary_material

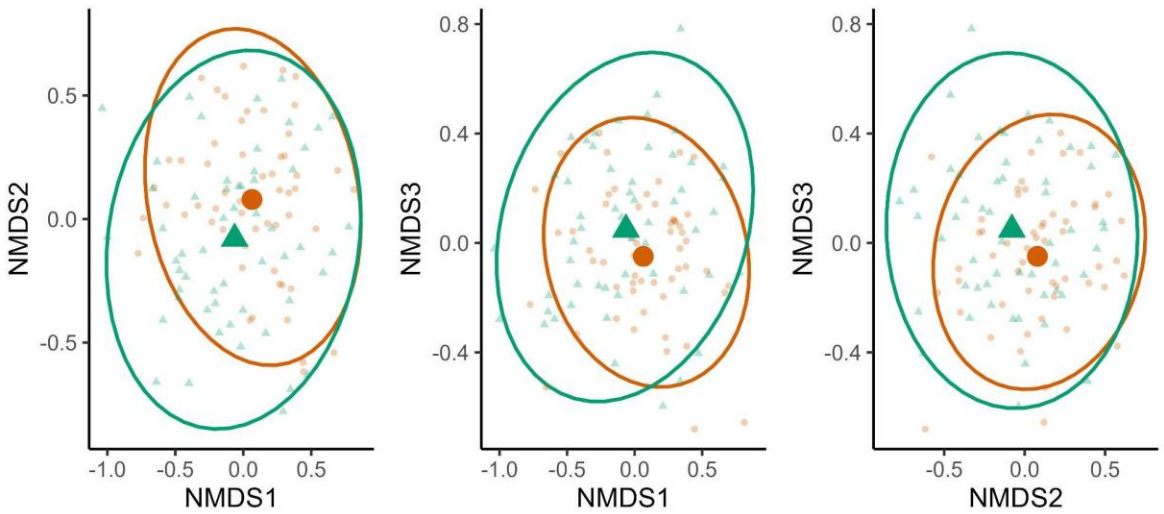


**Figure S1: Composition of plant communities in restored and control sections.**
Nonmetric multi-dimensional scaling (NMDS) with three-axis solution and stress = 0.169. Each small point and triangle represents a section, green triangles represent restored sections, orange dots are control sections.

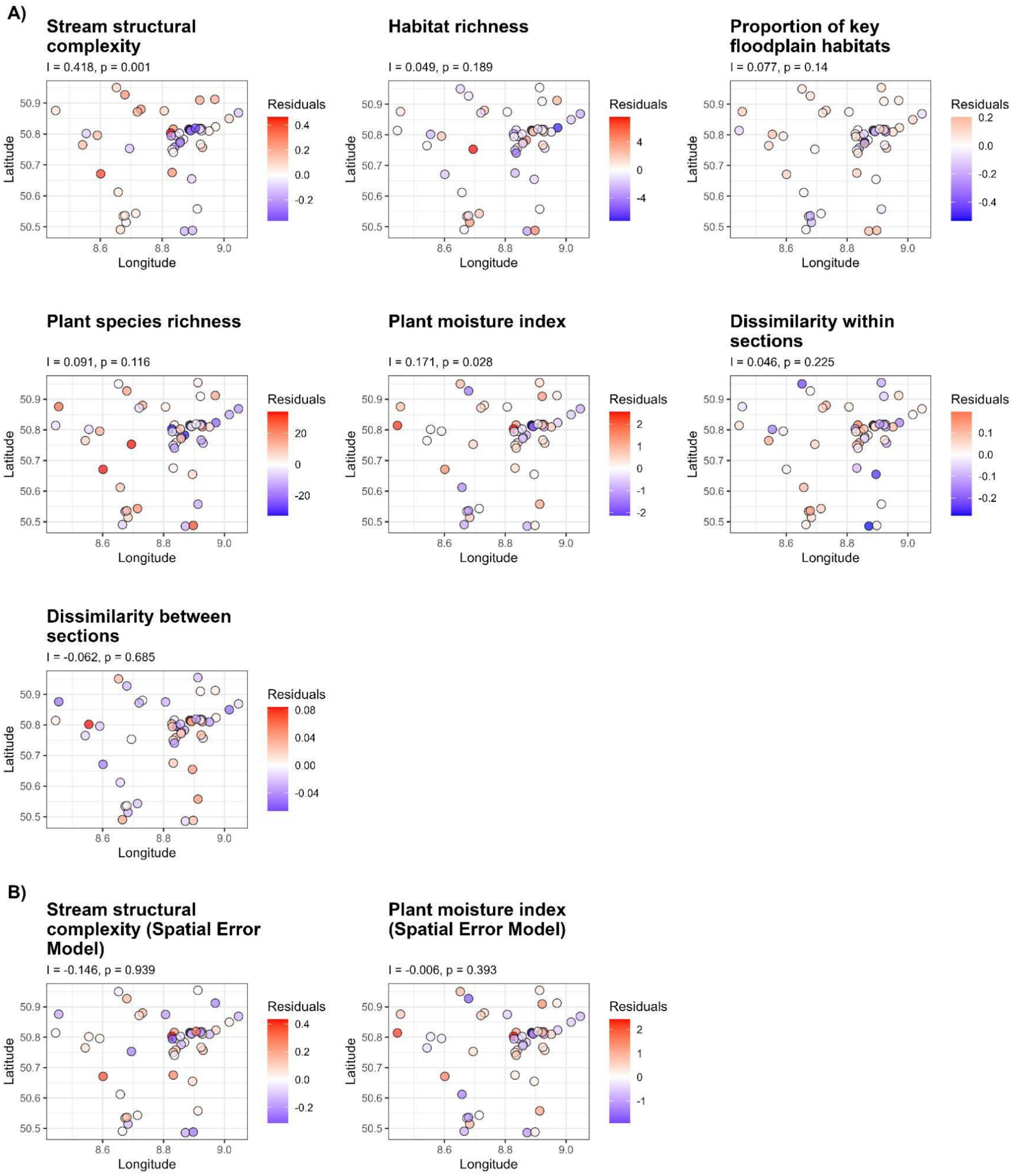


**Figure S2: Spatial distribution maps of residuals of each model used for the time since restoration analysis.**

**A)** Residuals from the linear models without accounting for spatial autocorrelation.

**B)** Residuals from models showing spatial autocorrelation, refitted using spatial error models.

**Table S3**: Full nomenclature of component models of the structural equation model.

| Model *type* | Response variable ~ | Explanatory variables + | | | | | Random factor | Specifications |
|---|---|---|---|---|---|---|---|---|
| Model 1 *lmer* | Structural complexity ~ | Area | | | | Site type | (1\|sitenr) | |
| Model 2 *lmer* | Habitat richness ~ | Area | | Structural complexity | | Site type | (1\|sitenr) | |
| Model 3 *glmmTMB* | Key floodplain habitats ~ | Area | Habitat richness | Structural complexity | | Site type | (1\|sitenr) | family = beta_family (link = "logit")) |
| Model 4 *lmer* | Species richness ~ | Area | Habitat richness | Structural complexity | | Site type | (1\|sitenr) | |
| Model 5 *lmer* | Moisture index ~ | Area | Habitat richness | Structural complexity | Key floodplain habitats | Site type | (1\|sitenr) | |
| Model 6 *lmer* | Dissimilarity between ~ | Area | Habitat richness | Structural complexity | | Site type | (1\|sitenr) | |

**Table S4: Habitat types in the stream sections studied.** We assigned the habitat types found in the study sections to their alliances and translated them into key-floodplain habitats A1 to A11 following (Januschke et al., 2024, P113). We made changes for three alliances that currently are not covered by this scheme but represent important habitats in the floodplains. These modifications are indicated with Asterisks in the code column.

| Habitats | Alliance | Code | | Key floodplain habitat |
|---|---|---|---|---|
| Acer platanoidis -Tilietum platyphylli | Tiolio platyphylli - Acerion pseudoplatani | | | |
| Aegopodio-Sambucetum nigrae | Balloto-Sambucion nigrae | | | |
| Aegopodium-Sambucetum nigrae | Balloto-Sambucion nigrae | | | |
| Alchemillo vulgaris-Arrhenatheretum elatioris | Arrhenatherion elatioris | 5.4.2.1. | A6 | Wet meadows |
| Angelico sylvestris-Scirpetum sylvatici | Calthion palustris | 5.4.1.5. | A6 | Wet meadows |
| Arctio-Artemisietum vulgaris | Arction lappae | 3.5.1.1. | A5 | Tall herb vegetation |
| Arrhenatheretum elatioris | Arrhenatherion elatioris | 5.4.2.1. | A6 | Wet meadows |
| Brometum sterilis | Sysymbrion officinalis | | | |
| Building | | | | |
| Calthion palustris | Calthion palustris | 5.4.1.5. | A6 | Wet meadows |
| Cardamine amarae-Petasitetum hybridi | Petasito hybridi-Chaerophyllion hirsuti | * | A5 | ıll herb vegetation |
| Caricetum acutiformis | Magnocaricion (syn. Caricion elatae) | 1.5.1.4. | A4 | Sedge and reedbeds |
| Caricetum distichae | Magnocaricion (syn. Caricion elatae) | 1.5.1.4. | A4 | Sedge and reedbeds |
| Caricetum versicariae | Magnocaricion (syn. Caricion elatae) | 1.5.1.4. | A4 | Sedge and reedbeds |
| Caricetum vulpinae | Magnocaricion (syn. Caricion elatae) | 1.5.1.4. | A4 | Sedge and reedbeds |
| Chaerophylletum aurei | Aegopodion podagrariae | 3.5.3.1. | A5 | Tall herb vegetation |
| Chaerophylletum aurei | Aegopodion podagrariae | 3.5.3.1. | A5 | Tall herb vegetation |

| | | | | |
|---|---|---|---|---|
| Chaerophylletum bulbosi | Aegopodion podagrariae | 3.5.3.1. | A5 | Tall herb vegetation |
| Cirsietum lanceolati-arvensis | Arction lappae | 3.5.1.1. | A5 | Tall herb vegetation |
| Convolvulo arvensis-Brometum inermis | Convolvulo-Agropyrion repentis | | | |
| Crataego-Prunetum spinosae | Carpino betuli-Prunion spinosae | | | |
| Cuscuto europea-Convolvuletum sepium | Senecion fluviatilis (syn. Convolvion sepii) | | A5 | Tall herb vegetation |
| Cynosuro cristati-Lolietum perennis | Cynosurion cristati | | | |
| Dactylido-Festucetum arundinaceae | Potentillion anserinae | | | |
| Eleocharitetum palustris | Eleocharito-Sagittarion sagittifoliae | * | A4 | Sedge and reedbeds |
| Epilobio hirsuti-Convolvuletum sepium | Senecion fluviatilis (syn. Convolvion sepii) | 3.5.2.2. | A5 | Tall herb vegetation |
| Epilobio hirsuti-Scrophularietum umbrosae | Glycerio-Sparganion emersi | 1.5.1.3. | A4 | Sedge and reedbeds |
| Epilobio-Juncetum effusi | Calthion palustris | 5.4.1.5. | A6 | Wet meadows |
| Equisetetum fluviatilis | Phragmition australis | 1.5.1.1. | A4 | Sedge and reedbeds |
| Fallow land | | | | |
| Festuco rubrae-Cynosuretum cristati | Cynosurion cristati | | | |
| Field | | | | |
| Filipendulo ulmariae-Geranietum palustris | Filipendulion ulmariae | 5.4.1.2. | A5 | Tall herb vegetation |
| Fixed streambed | | | | |
| Flowering strip | | | | |
| Frangulo-Salicetum auritae | Salicion cinereae | 8.2.1.2 | A9 | Carr |
| Frangulo-Salicetum cinereae | Salicion cinereae | 8.2.1.2 | A9 | Carr |
| Fruit shrub | | | | |
| Galio molluginis-Alopecuretum pratensis | Arrhenatherion elatioris | 5.4.2.1. | A6 | Wet meadows |
| Galio palustris-Caricetum ripariae | Magnocaricion (syn. Caricion elatae) | 1.5.1.4. | A4 | Sedge and reedbeds |
| Gravel bank | | | | |
| Impatienti glanduliferae-Convolvuletum sepium | Senecion fluviatilis (syn. Convolvion sepii) | 3.5.2.2. | A5 | Tall herb vegetation |
| Juncetum acutiflori | Calthion palustre | 5.4.1.5. | A6 | Wet meadows |
| Lolietum perennis | Cynosurion cristati | | | |

| | | | | |
|---|---|---|---|---|
| Loto uliginosi-Holcetum lanati | Calthion palustris | 5.4.1.5. | A6 | Wet meadows |
| Mud bank | | | | |
| Pasture | | | | |
| Path | | | | |
| Phalaridetum arundinaceae | Magnocaricion (syn. Caricion elatae) | 1.5.1.4. | A4 | Sedge and reedbeds |
| Phalarido arundinaceae-Petasitetum hybridi | Aegopodion podagrariae | 3.5.3.1. | A5 | Tall herb vegetation |
| Phragmitetum australis | Phragmition australis | 1.5.1.1. | A4 | Sedge and reedbeds |
| Piping | | | | |
| Plantagini lanceolatae-Festucetum rubrae | Arrhenatherion elatioris | 5.4.2.1. | A6 | Wet meadows |
| Poo trivialis-Rumicetum obstusifolii | Potentillion anserinae | | | |
| Poo-Trisetum flavescentis | Arrhenatherion elatioris | 5.4.2.1. | A6 | Wet meadows |
| Potentilla anserina-community | Potentillion anserinae | | | |
| Pruno-Fraxinetum | Alno-Ulmion | 8.4.3.3. | A1 0 | Hardwood riparian forest |
| Pruno-Ligustretum | Berberidion | | | |
| Prunus domestica-community | Balloto-Sambucion nigrae | | | |
| Ranunculo repentis-Alopecuretum geniculati | Potentillion anserinae | * | A6 | Wet meadows |
| Ranunculus repens-community | Potentillion anserinae | | | |
| Riparian bank (unvegetated) | | | | |
| Rorippo sylvestris-Phalaridetum arundinaceae | Glycerio-Sparganion emersi | 1.5.1.3. | A4 | Sedge and reedbeds |
| Row of trees | | | | |
| Rubetum idaei | Sambuco racemosae-Salicion capreae | | | |
| Rubo fructicosi-Coryletum avellanae | Carpino betuli-Prunion spinosae | | | |
| Rubo-Calamagrostietum epigeji | Convolvulo-Agropyrion repentis | | | |
| Rubus spec.-shrub | | | | |
| Rumici-Alopecuretum aequalis | Bidention tripartiae | 3.2.1.1. | A3 | Banks with little or no vegetation |
| Salicetum albae | Salicion albae | 8.1.1.2 | A8 | Softwood riparian forest |

| | | | | |
|---|---|---|---|---|
| Salicetum caprae | Sambuco racemosae-Salicion capreae | | | |
| Salicetum elagni-purpureae | Salicion eleagni (Salicion eleagno-daphnoidis) | 8.1.1.1. | A8 | Softwood riparian forest |
| Salicetum fragilis | Salicion albae | 8.1.1.2 | A8 | Softwood riparian forest |
| Salicetum triandrae | Salicion albae | 8.1.1.2 | A8 | Softwood riparian forest |
| Salix purpurea- community | Salicion albae | 8.1.1.2 | A8 | Softwood riparian forest |
| Seeded grassland | | | | |
| Slope | | | | |
| Sparganio-Glycerietum fluitantis | Glycerio-Sparganion emersi | 1.5.1.3. | A4 | Sedge and reedbeds |
| Stellario holosteae-Carpinetum betuli | | | | |
| Street | | | | |
| Tanacetum vulgaris-Arrhenatheretum elatioris | Arrhenatherion elatioris | 5.4.2.1. | A6 | Wet meadows |
| Typhetum latifoliae | Phragmition australis | 1.5.1.1. | A4 | Sedge and reedbeds |
| Urtico dioicae-Aegopodietum podagrariae | Aegopodion podagrariae | 3.5.3.1. | A5 | Tall herb vegetation |
| Urtico-Alnetum glutinosae | Alnion glutinosae | 8.2.1.1. | A9 | Carr |
| Urtico-Salicetum cineraea | Salicion cinereae | 8.2.1.2 | A9 | Carr |
| Valeriano officinalis-Filipenduletum ulmariae | Filipendulion ulmariae | 5.4.1.2. | A5 | Tall herb vegetation |
| Water | | | | |

**S5:** Summary of the sampling process for stream structural complexity and explanation of modifications.

To assess the stream structural complexity, we followed the methodology described by the German Working Group on Water Issues (LAWA, 1999). The protocol evaluates the complexity of 100 m stream stretches using six parameters: (1) complexity of flow, (2) longitudinal profile, (3) cross profile, (4) riverbed structure, (5) shore structure, and (6) surroundings. Each parameter includes several sub-parameters that are assessed in the field and assigned to predefined categories. Each selectable category then refers for the calculation to a value between 1 and 7, where 7 is the worst (lowest complexity) and 1 is the best category (highest complexity). For each parameter, sub-parameter scores are averaged to obtain a parameter-specific value. The overall stream structural complexity index of the 100 m stream stretch is then calculated as the mean of the six parameter values.

*Modification applied in this study*

For the analyses presented in the present paper, we excluded parameter 6 (“Surroundings”) from the calculation of stream structural complexity. We used structural complexity as an indicator of the intensity of in-stream restoration measures and to test its effect on plant diversity and habitat diversity within the project area. Including the “Surroundings” parameter would have introduced conceptual overlap between predictor and response variables because this parameter directly reflects characteristics of the adjacent vegetation and habitat structure that are also captured by the response variables. Its inclusion would increase the risk of inflated explained variance. We therefore only used parameters 1 to 5 to calculate the respective stream structural complexity value of each 100 m stretch and averaged the values of all stretches within a section to obtain one stream structural complexity index value for each section. For a more easy and intuitive interpretation of the stream structural complexity index, we rescaled and inverted the original index value, so that the index now ranges between 0 and 1, where zero is the worst stream structural complexity and 1 is the highest.
